# Long-read sequencing reveals novel mitochondrial genome variants undetected by short-read sequencing in Korean population

**DOI:** 10.64898/2026.07.15.738850

**Authors:** Hyung Jun Kim, Sung Min Kim, Kyungheon Yoon, Bong-Jo Kim, Han Jun Jin, Young Jin Kim

## Abstract

While long-read sequencing technologies (e.g., PacBio Revio, ONT) have revolutionized high-quality genome assembly for the human pangenome, mitochondrial genome (mtDNA) analysis still largely relies on short-read and Sanger sequencing. However, short-read sequencing often lacks the resolution required to resolve complex variations due to the unique features of mtDNA, such as high mutation rates and repetitive homopolymeric regions, which frequently lead to alignment artifacts and mapping ambiguities. To address this, we evaluated whether applying long-read sequencing to mtDNA improves analytical quality in empirical data. Through comprehensive bioinformatics analyses, we compared the performance of long-read sequencing against short-read sequencing and microarrays. Our results revealed that long-read sequencing detected the highest number of variants (n = 533), significantly outperforming both short-read sequencing (n = 525) and microarrays (n = 49). Notably, both sequencing methods provided significantly higher resolution in haplogroup assignment compared to microarrays in terms of phylogenetic depth (*p* < 0.05). Long-read sequencing demonstrated superior detection power, particularly for InDels. We identified two novel non-synonymous variants, including a unique InDel detected exclusively by long-read sequencing. Protein modeling and stability analysis validated that this InDel causes structural instability (RMSD > 2.0 Å, *ΔΔ*G = -45.21 kcal/mol). Furthermore, we confirmed that this novel InDel is shared among haplogroup A samples in both the 1000 Genomes Project ONT dataset and the Korean population, highlighting the practical implications of long-read sequencing for molecular biology and population genetics.

## Introduction

The mitochondrial genome (mtDNA) occupies a unique position in human genetics, serving simultaneously as a functional component of cellular energy production and as a convenient molecular marker for tracing human evolutionary history (da Fonseca et al. 2008; St. John 2025). By encoding 13 essential subunits of the oxidative phosphorylation (OXPHOS) complexes, mtDNA plays an essential role in bioenergetics and cellular metabolism (Gilkerson et al. 2013; Ferreira and Rodriguez 2024). Equally, its strictly maternal inheritance, lack of recombination, and elevated mutation rate have rendered mtDNA an indispensable tool for reconstructing ancient migration routes and characterizing population-level genetic diversity (Stewart and Chinnery 2015). Despite this dual significance, the genomic architecture of mtDNA remains incompletely resolved, in large part because of analytical challenges intrinsic to the mitochondrial system itself (Laricchia et al. 2022; Macken et al. 2023; Slapnik et al. 2024).

These analytical challenges stem directly from fundamental differences between mitochondrial and nuclear genetics (Ferreira and Rodriguez 2024). As a relic of its endosymbiotic origin, the mitochondrial genome retains a compact, circular structure (Sharma and Sampath 2019). Residing in the mitochondrial matrix, mtDNA is chronically exposed to reactive oxygen species (ROS) produced during electron transport, and its replication occurs independently of the cell cycle—resulting in elevated mutation rates and extensive copy number variability (heteroplasmy) within and between cells (Stewart and Chinnery 2015; Ferreira and Rodriguez 2024). These biological properties translate into concrete computational challenges: alignment algorithms must accommodate a circular topology, variant callers must quantify heteroplasmic allele fractions across thousands of copies, and pipelines must distinguish true mitochondrial variants from nuclear-embedded mitochondrial sequences (NUMTs) (Xue et al. 2023). Standard bioinformatics workflows designed for the diploid, linear nuclear genome do not natively address these requirements (Laricchia et al. 2022).

For over a decade, Short-Read Sequencing (SRS) platforms such as Illumina have served as the dominant technology for genomic analysis (Macken et al. 2023). Yet when applied to mitochondrial DNA, SRS encounters two categories of limitation that compromise variant detection accuracy. The first is technical. Short reads (typically 150 bp) cannot span the repetitive and homopolymeric regions abundant in the mitochondrial genome, leading to ambiguous alignments. More critically, the nuclear genome harbors thousands of nuclear mitochondrial DNA segments (NUMTs)—pseudogene copies of mtDNA integrated into nuclear chromosomes—that share high sequence identity with their mitochondrial counterparts (Wei et al. 2022). Short reads originating from NUMTs are frequently misaligned to the mitochondrial reference, inflating false-positive variant calls (Xue et al. 2023). The second limitation is conceptual. Conventional pipelines align reads to the revised Cambridge Reference Sequence (rCRS), a single linear representation derived from a European haplogroup H individual (Andrews et al. 1999). This design inherently introduces reference bias: insertions, deletions, and highly divergent alleles not represented in rCRS are systematically underdetected (Zaragoza et al. 2011; Rubin et al. 2023). Consequently, population-scale databases have historically filtered variants in “noise-prone” regions, potentially discarding signals of genuine biological significance (Laricchia et al. 2022).

The emergence of Long-Read Sequencing (LRS) technologies offers a path to overcoming both categories of SRS limitation. In particular, PacBio high-fidelity (HiFi) sequencing generates reads exceeding 10 kb with per-base accuracy up to 99.9%, enabling full-length coverage of the ∼16.6 kb mitochondrial genome in single continuous molecules (Wenger et al. 2019). This capability eliminates fragmentation-induced alignment ambiguity and permits unambiguous discrimination of true mitochondrial variants from NUMT-derived artifacts (Wei et al. 2022; Macken et al. 2023). At the reference level, the success of graph-based pangenomes for the nuclear genome—as demonstrated by the Human Pangenome Reference Consortium (HPRC)—has established both the conceptual framework and computational infrastructure for moving beyond single linear references (Liao et al. 2023). Extending this paradigm to the mitochondrial genome is a logical next step: a mitochondrial pangenome would capture population-specific structural variants and haplogroup-defining alleles that rCRS-based analyses systematically miss. However, constructing such a resource requires high-quality, long-read-resolved mitochondrial assemblies from diverse populations—precisely the data that remain scarce for non-European cohorts.

The Korean population represents one such critically underrepresented group. In gnomAD v3.1, East Asian samples constitute only ∼3% of the mitochondrial dataset, with Korean individuals comprising a small fraction thereof (Laricchia et al. 2022). This underrepresentation is consequential: the Korean population harbors a distinctive mitochondrial haplogroup distribution—enriched in macro-haplogroups D4, M7, and B, with frequencies and sub-lineage structures that diverge substantially from neighboring Chinese and Japanese populations (Lee et al. 2006; Jin et al. 2009). Without adequate representation in reference databases and pangenomic resources, population-specific variants risk being misclassified as noise or excluded from clinical interpretation.

Here, we expanded our understanding of mitochondrial genome using long-read sequencing on 65 Korean individuals. Using the high-quality variant calls from LRS, we characterize the mitochondrial genomic landscape of the Korean population, including haplogroup distribution, population structure, and phylogenetic relationships within an East Asian context. Additionally, we present a systematic cross-platform evaluation of mtDNA variant detection in a Korean population cohort, comparing Long-Read Sequencing (PacBio Revio HiFi) with widely used platforms such as Short-Read Sequencing (Illumina NovaSeq) and genotyping microarray (Korea Biobank Array). By benchmarking these three technologies against a common sample set, we establish the analytical resolution achievable by each platform and identify variant classes—particularly homopolymeric insertions/deletions and low-frequency heteroplasmies—for which LRS provides measurable improvements in sensitivity and specificity. Together, these analyses provide both a methodological framework for mitochondrial variant detection and foundational genomic resources toward the future construction of a mitochondrial pangenome for East Asian populations.

## Material & methods

### Study Population and Data Resources

This study analyzed whole mitochondrial genome sequences from 65 Korean individuals. Participants were randomly selected from the Korean Genome and Epidemiology Study (KoGES) cohort, a large-scale population-based prospective study comprising community-based (Ansan-Ansung), urban-based, and rural-based sub-cohorts designed to investigate the genetic and environmental determinants of common complex diseases in the Korean population (Kim et al. 2017). All participants provided written informed consent, and the study protocols were reviewed and approved by the Institutional Review Board of the Korea Disease Control and Prevention Agency (KDCA) (Approval No. KDCA-2024-02-09).

To facilitate cross-platform validation and comparative population genetic analyses, we incorporated two external high-coverage mitochondrial genome datasets. First, the 1000 Genomes Project (1KGP) short-read dataset, which comprises 2,503 unrelated individuals sequenced to ∼30× coverage on the Illumina platform using GRCh38 as the reference genome (Byrska-Bishop et al. 2022). Second, the Vienna Oxford Nanopore Technologies (ONT) long-read dataset from the 1KGP structural variation study, which includes 1,019 individuals sequenced by long-read technology (Schloissnig et al. 2025). These datasets were used for orthogonal variant validation and cross-population haplogroup comparison (**Supplementary Table S1**).

### Sequencing and Genotyping Procedures

To evaluate platform-specific variant detection performance, genomic data from the 65 Korean and 1KGP individuals were generated using three distinct platforms: long-read sequencing (LRS), short-read sequencing (SRS), and microarray genotyping.

### Long-read Sequencing (PacBio Revio)

High-molecular-weight (HMW) genomic DNA was extracted from peripheral whole blood using standard salting-out methods. DNA quality and concentration were assessed using a Nanodrop spectrophotometer (Thermo Fisher Scientific) and Qubit fluorometer (Invitrogen). For library preparation, approximately 5 µg of HMW gDNA was sheared to a target size of 15–18 kb using the Megaruptor 3 system (Diagenode). SMRTbell libraries were constructed using the SMRTbell Prep Kit 3.0 (Pacific Biosciences), following the manufacturer’s standard protocol. This process involved DNA damage repair, end-repair/A-tailing, and ligation of overhang adapter v3. Non-SMRTbell molecules were removed by nuclease treatment, and size selection was performed to enrich for fragments >10 kb. The final libraries were sequenced on the PacBio Revio system using Revio SMRT Cells, generating high-fidelity (HiFi) circular consensus sequencing (CCS) reads with a mean per-base accuracy of Q30+ (Wenger et al. 2019). The mean sequencing depth for mitochondrial reads was 254.81× (**Table 1**).

**Table 1.** Summary of mitochondrial genome variants by sequencing platforms.

| Sequencing Platform | Variant Type | Number of variants |  |  | Mean TLOD score | Mean Depth |
| --- | --- | --- | --- | --- | --- | --- |
|  |  | Total | SNVs | InDels |  |  |
| Longread<br>(n = 65) | Known | 531 | 503 | 28 | 1006.21 | 254.81 |
|  | Novel | 2 | 1 | 1 | 54.33 | 195.00 |
| Shortread<br>(n = 65) | Known | 524 | 503 | 21 | 12144.54 | 3530.66 |
|  | Novel | 1 | 1 | 0 | 1636.04 | 3930.00 |
| KBAv1.1*<br>(n = 65) | Known | 54 | 54 | 0 | NA | NA |
|  | Novel | 0 | 0 | 0 |  |  |

\*Among the 180 SNVs, 54 were identified as non-monomorphic
Abbreviations: SNVs, single nucleotide variants; InDels, insertion/deletion; TLOD, tumor log odds; KBAv1.1, Korea biobank array version 1.1

### Short-read Sequencing (Illumina)

Short-read sequencing was performed following the whole-genome sequencing (WGS) workflow described by (Hwang et al. 2022). Briefly, 1 µg of genomic DNA was quantified using the Quant-iT PicoGreen dsDNA Assay Kit (Invitrogen) and fragmented to an average size of 350 bp using adaptive focused acoustics (Covaris). Sequencing libraries were prepared using the TruSeq DNA PCR-Free Library Prep Kit (Illumina) to minimize amplification bias. The libraries underwent end-repair, size selection, and adenylation, followed by adapter ligation. Quality control was performed using an Agilent 2100 Bioanalyzer. Sequencing was conducted on the Illumina NovaSeq 6000 platform, generating 150 bp paired-end reads to a target depth of ∼30× for nuclear genome coverage, which yielded a substantially higher mitochondrial read depth (mean depth: 3,530.63×) due to the high copy number of mtDNA per cell (**Table 1**).

### Microarray Genotyping

Large-scale genotyping was performed using the Korea Biobank Array (KBA) v1.1, a population-optimized SNP array designed specifically for the Korean population, containing approximately 830,000 markers including 180 mitochondrial SNP probes (Moon et al. 2019). Total genomic DNA (200 ng) was amplified, fragmented into 25–125 bp segments, and hybridized to the array plates for 24 hours at 48°C. Following hybridization, the arrays were washed and stained using the GeneTitan Fluidics Station 450 and scanned on the GeneTitan Multi-Channel Instrument (Thermo Fisher Scientific).

### Sequence Alignment and Mitochondrial Read Extraction

#### Short-Read Sequencing (Illumina NovaSeq) and LRS Mitochondrial Read Processing

Short-read whole-genome sequencing data were generated following the workflow described by Hwang et al. (2022). Briefly, raw FASTQ reads were preprocessed using fastp (v0.23.2) for adapter trimming and quality filtering (Phred score < 20), then aligned to GRCh38 using BWA-MEM (v0.7.17) with default parameters. The mean sequencing depth for mitochondrial reads was 3,530.63× (**Table 1**).

For mitochondrial-specific variant calling, reads mapping to chrM were extracted from the whole-genome BAM using GATK PrintReads with --read-filter MateOnSameContigOrNoMappedMateReadFilter and --read-filter MateUnmappedAndUnmappedReadFilter. Extracted reads were reverted to unaligned format using GATK RevertSam, converted to interleaved FASTQ using GATK SamToFastq, and realigned to the GRCh38 mitochondrial reference (rCRS; NC_012920.1) using BWA-MEM (v0.7.17) with the -p flag for interleaved paired-end input. Aligned and unaligned BAMs were merged using GATK MergeBamAlignment with --MAX_INSERTIONS_OR_DELETIONS -1. Duplicate reads were marked using GATK MarkDuplicates with --OPTICAL_DUPLICATE_PIXEL_DISTANCE 2500 and sorted by coordinate using GATK SortSam.

An identical pipeline was applied to the PacBio HiFi long-read data for mitochondrial variant calling, with the same GATK workflow (PrintReads → RevertSam → SamToFastq → BWA-MEM → MergeBamAlignment → MarkDuplicates → SortSam) to ensure methodological consistency across the two sequencing platforms.

#### ONT Long-Read Data Processing

For the 1KGP ONT long-read dataset (n = 1,019; Schloissnig et al., 2025), mitochondrial reads were extracted from the provided BAM files and processed through a customized GATK mitochondrial variant calling pipeline. Read groups were assigned using GATK AddOrReplaceReadGroups with platform designation set to ONT. Reads were reverted to unaligned format using GATK RevertSam, then realigned to the GRCh38 mitochondrial reference using BWA-MEM (v0.7.17) with the -x ont2d flag to optimize alignment for Oxford Nanopore two-directional reads. Aligned and unaligned BAMs were merged using GATK MergeBamAlignment with --MAX_INSERTIONS_OR_DELETIONS -1 to accommodate the higher InDel rate characteristic of nanopore sequencing.

Because a subset of ONT reads lacked base quality scores (represented as ‘*’ in the BAM QUAL field), a quality score injection step was applied prior to variant calling: reads with missing quality strings were assigned a uniform Phred score of 37 (ASCII ‘F’) across all bases using an AWK-based in-line filter to satisfy GATK input requirements. Secondary and supplementary alignments were excluded (SAMtools flag -F 2304). Duplicate reads were marked using GATK MarkDuplicates and sorted by coordinate using GATK SortSam.

#### Mitochondrial Variant Calling

Mitochondrial variants from LRS, SRS, and 1KGP ONT data were identified using GATK (v4.4.0) Mutect2 in mitochondria mode, following the GATK Best Practices pipeline for mitochondrial variant calling (Laricchia et al. 2022). This pipeline addresses the unique analytical challenges of mtDNA, including its circular topology and the potential for nuclear mitochondrial DNA segments (NUMTs) to confound variant calls.

Specifically, the workflow involved: (i) separate variant calling on the non-control region (chrM:576–16024) using the standard rCRS reference and on the control region using a shifted reference (origin at position 8,000; calling interval chrM:8025–9144 on the shifted coordinate) to resolve artifacts at the D-loop junction; (ii) variant calling with Mutect2 using --mitochondria-mode true, --max-reads-per-alignment-start 75, and --max-mnp-distance 0; (iii) merging of control-region calls back to the standard coordinate system using GATK LiftoverVcf, followed by GATK MergeVcfs to combine coding and control region calls.

For the 1KGP ONT data, two additional Mutect2 flags were applied to accommodate nanopore-specific read characteristics: --disable-read-filter WellformedReadFilter (to accept reads with injected quality scores) and --dont-use-soft-clipped-bases true (to exclude soft-clipped bases from variant calling, reducing ONT-specific alignment edge artifacts). These flags were not required for the Korean cohort LRS and SRS data, which used standard Mutect2 parameters.

Variants were filtered using GATK FilterMutectCalls with --mitochondria-mode, --max-alt-allele-count 4, and --min-allele-fraction 0. Known artifact sites were masked using GATK VariantFiltration with a blacklist BED file (Laricchia et al. 2022). Multi-allelic sites were decomposed using GATK LeftAlignAndTrimVariants with --split-multi-allelics and --dont-trim-alleles, and only PASS-filtered variants were retained using GATK SelectVariants.

For both the Korean cohort and the 1KGP ONT data, mitochondrial contamination was estimated using haplocheck (Weissensteiner et al. 2020). When haplocheck flagged a sample as contaminated, the contamination level was calculated as 1 minus the major haplogroup heteroplasmy fraction; otherwise, contamination was set to zero. This estimate was supplied to a second round of FilterMutectCalls via the --contamination-estimate parameter. Variants were then re-filtered with the contamination-adjusted threshold and re-masked against the blacklist before final output.

#### VCF Normalization and Variant Harmonization

To ensure consistent variant representation across platforms and enable accurate cross-platform concordance analysis, VCF files from all sequencing platforms were processed through a custom normalization pipeline.

First, multi-allelic variants were decomposed and left-aligned against the rCRS reference using bcftools norm (v1.17) with the -m - flag.

Second, each variant was annotated with its HGVS mitochondrial nomenclature (e.g., NC_012920.1:m.1234A>G) using the HGVS Python library (v1.5.4) connected to a local Universal Transcript Archive (UTA) database, enabling standardized variant identification independent of VCF positional representation.

Third, genotype (GT) fields were corrected for mitochondrial haploidy: heterozygous calls (0/1) with variant allele frequency (VAF) ≥ 0.90 were reclassified as homozygous alternative (1/1), reflecting the expectation that high-frequency mitochondrial alleles represent near-homoplasmic states rather than true heterozygosity in the nuclear diploid sense. Calls with intermediate VAF (0.10 ≤ VAF < 0.90) were retained as heterozygous (0/1) to preserve heteroplasmic variants, whereas calls with VAF < 0.10 were reclassified as homozygous reference (0/0), as such low-frequency signals likely reflect sequencing noise rather than biologically meaningful heteroplasmy.

Fourth, VCF records were re-normalized to match database-standard representations at known problematic loci. This included: (a) resolution of poly-C tract insertions at positions 309, 573, 960, 5899, and 8278, where multiple VCF records representing the same underlying insertion were merged into a single representative record; (b) correction of D-loop hypervariable region variants near position 16,189, where GATK Mutect2 outputs context-dependent deletion representations that were converted to individual deletion records at canonical positions (e.g., m.16193del, m.16187_16188del) based on a priority-ordered rule set; (c) splitting of the compound deletion m.513_515del into its constituent components (m.513_514del and m.515del); and (d) re-anchoring of the CCCCCTCTA insertion at position 309 (originally called as m.315_316insCCCCCTCTA) to the database-standard representation at m.309_310ins.

#### Microarray Variant Calling and Quality Control

Large-scale genotyping was performed using the Korea Biobank Array (KBA) v1.1, a population-optimized SNP array containing approximately 830,000 markers including 180 mitochondrial SNP probes (Moon et al. 2019). Genotypes were called using the Axiom Analysis Suite with strict quality control thresholds (Dish QC ≥ 0.82 and Call Rate ≥ 97%).

For mitochondrial variants, genotype cluster plots for all 180 mitochondrial SNP probes were individually inspected using the Axiom Analysis Suite to verify genotype calling accuracy. Probes were excluded if they exhibited ambiguous cluster separation between genotype groups, showed evidence of batch effects, or had high missing rates across samples. This manual curation was performed to mitigate the known limitation that automated genotype calling algorithms optimized for diploid autosomal loci may produce unreliable calls for haploid mitochondrial markers, particularly at positions with rare alleles or low-frequency heteroplasmy. The 54 non-monomorphic variants passing quality control were exported in VCF format for cross-platform comparison.

#### Variant Annotation and Classification

Identified variants from all three platforms were annotated using the Ensembl Variant Effect Predictor (VEP, v110) (McLaren et al. 2016). To assess population frequencies and potential clinical significance, variants were cross-referenced against public databases including gnomAD v3.0 (Laricchia et al. 2022), HelixMTdb, NCBI dbSNP, and MITOMAP. Protein-level pathogenicity for missense variants was further evaluated using AlphaMissense (Cheng et al. 2023). Variants were classified as “novel” if absent from all four databases (gnomAD, MITOMAP, HelixMTdb, and NCBI).

#### Haplogroup Assignment and Population Genetic Analysis

Mitochondrial haplogroups were assigned for all 65 individuals across the three platforms using HaploGrep3 based on the PhyloTree Build 17 (Forensic Update 1.2) classification (van Oven and Kayser 2009; Schönherr et al. 2023). Haplogroup assignment accuracy was evaluated by comparing the phylogenetic depth (the number of hierarchical levels resolved), the HaploGrep3 quality score, and the number of expected variants missing for each assigned haplogroup across platforms. Statistical differences were assessed using paired t-tests with LRS as the reference platform.

Principal Component Analysis (PCA) was performed to assess the population genetic structure of the 65 Korean individuals in the context of global diversity. Mitochondrial variant calls from the Korean cohort were merged with those from the 1KGP Phase 3 dataset (comprising African [AFR], European [EUR], South Asian [SAS], East Asian [EAS], and admixed American [AMR] super-populations). PCA was conducted separately for LRS-derived and SRS-derived variant sets to assess platform consistency in capturing population structure.

#### Protein Structure Modeling and Molecular Dynamics Simulations

To assess the structural and functional consequences of novel variants, we performed in silico protein modeling and molecular dynamics (MD) simulations for two variants of interest: the *MT-ND1* frameshift variant (m.4247 TT>T; p.Ile314X) and the *MT-CYB* missense variant (m.15635 T>A; p.Ser297Thr).

The cryo-EM structure of the human mitochondrial respiratory Complex I (NADH:ubiquinone oxidoreductase; PDB ID: 5XTD) served as the wild-type (WT) template for ND1 protein, and the crystal structure of cytochrome bc1 complex (PDB ID: 1PPJ) was used for CYTB protein. Mutant structures were modeled using AlphaFold3 to predict conformational changes induced by the variants(Abramson et al. 2024). Structural visualization and superimposition were performed using UCSF ChimeraX (Pettersen et al. 2021). Intermolecular interaction networks, including hydrogen bonds and hydrophobic contacts, were analyzed using LigPlot+ to characterize changes in local binding environments (Laskowski and Swindells 2011).

Thermodynamic stability changes upon mutation (ΔΔG) were estimated using FoldX (v5.0) (Schymkowitz et al. 2005). The wild-type structure was first energy-minimized using the FoldX RepairPDB module, and ΔΔG values were calculated as the difference in folding free energy between mutant and wild-type conformations (ΔΔG = ΔG_mutant – ΔG_wildtype).

For MD simulations, the ND1 wild-type and mutant (p.Ile314X) structures were embedded in a lipid bilayer mimicking the mitochondrial inner membrane using the CHARMM-GUI Membrane Builder (Wu et al. 2014). The membrane composition was set to a phosphatidylcholine-to-phosphatidylethanolamine (PC:PE) ratio of 40:35, consistent with established mitochondrial membrane lipid compositions (Horvath and Daum 2013). The systems were solvated with TIP3P water model (Jorgensen et al. 1983). To mimic the physiological environment of mitochondria, the systems were neutralized by adding KCl counterions to a final concentration of 150 mM (Garlid and Paucek 2003). We employed the CHARMM36m force field for proteins and lipids. Hydrogen Mass Repartitioning (HMR) was applied to enable a 4 fs integration time step. After energy minimization and equilibration (NVT for 500 ps, NPT for 1 ns), production MD simulations were performed for 100 ns using GROMACS (v2023.3) (Abraham et al. 2015). Trajectory analyses including Root Mean Square Deviation (RMSD), Root Mean Square Fluctuation (RMSF), Radius of Gyration (Rg), Solvent Accessible Surface Area (SASA), and hydrogen bond counts were computed using GROMACS built-in analysis modules.

#### Cross-Platform Variant Validation

To evaluate cross-platform concordance, variants detected by all three platforms were compared using a Venn diagram approach. For the sequencing platforms (LRS and SRS), variant allele frequencies (VAFs) were directly compared for genotype mismatched regions. Allele frequency correlations between the Korean cohort and gnomAD v3.0 East Asian (EAS) frequencies were assessed using Pearson’s correlation coefficient.

#### Statistical Analysis

Statistical analyses were performed using Python (v3.10) and R (v4.3.1). Paired t-tests were used to compare haplogroup assignment depth, quality scores, and missing variant counts between platforms, with LRS as the reference. Pearson’s correlation coefficients were calculated to assess the concordance of variant allele frequencies across platforms and against gnomAD v3.0 EAS population frequencies. Data visualizations were generated using matplotlib (v3.8) and seaborn (v0.13) in Python, and ggplot2 (v3.4) in R. Venn diagrams were generated using the VennDiagram R package. All statistical tests were two-sided, and p < 0.05 was considered statistically significant.

## Results

### Long-Read Sequencing Reveals an Expanded Mitochondrial Variant Landscape

Long-read sequencing (LRS) of whole mitochondrial genomes from 65 Korean individuals using PacBio Revio HiFi (mean depth: 254.81×; **Table 1**) identified 533 variants comprising 504 single-nucleotide variants (SNVs) and 29 insertions/deletions (InDels)—the most comprehensive variant catalog among the three platforms evaluated. Of these, 531 were classified as “known” (present in at least one of four public databases: gnomAD v3.0, MITOMAP, HelixMTdb, or NCBI dbSNP), while two were novel (absent from all four): a missense SNV at m.15635 T>A (p.Ser297Thr) in *MT-CYB* (VAF = 0.129, QUAL = 76.24) and a frameshift deletion in *MT-ND1* at m.4247 TT>T (VAF = 0.103, QUAL = 32.41). All 533 variants passed the VAF ≥ 0.10 heteroplasmy threshold used throughout the primary analyses in this study.

The LRS-derived variant set demonstrated strong concordance with established population databases. Allele frequencies of the 533 known variants showed a Pearson correlation coefficient of 0.988 with gnomAD v3.0 East Asian (EAS) population frequencies (**Figure 2b**), confirming that the cohort-level frequency spectrum captured by LRS faithfully recapitulates the expected East Asian mitochondrial allele frequency distribution. Similar agreement was observed with HelixMTdb (r = 0.872) and MITOMAP Polymorphism (r = 0.946) frequencies, although these databases reflect global rather than East Asian–specific frequencies and therefore yield lower nominal correlations.

**Figure 1.**
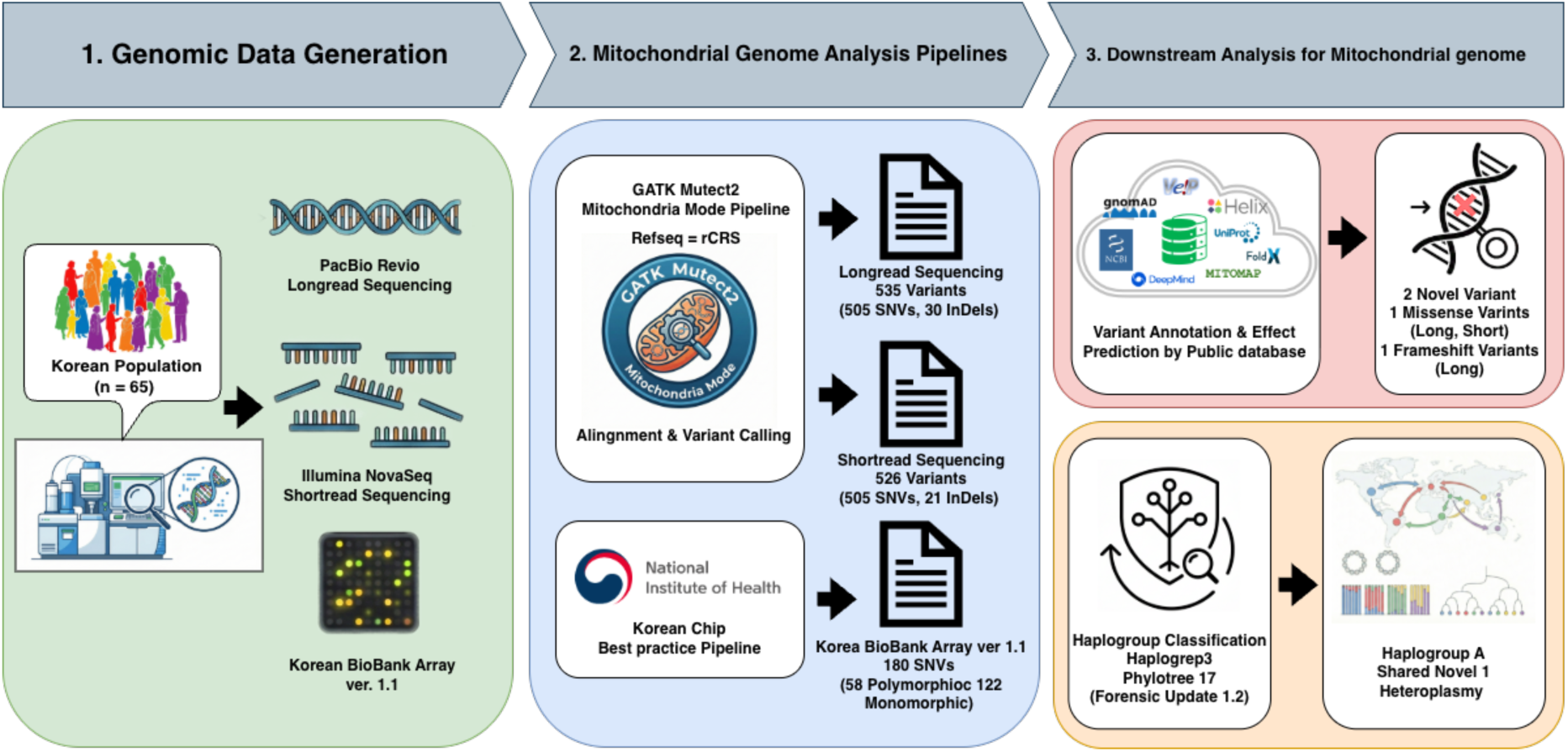
Overview of Comparative evaluation of sequencing platforms for mitochondrial genome analysis

**Figure 2.**
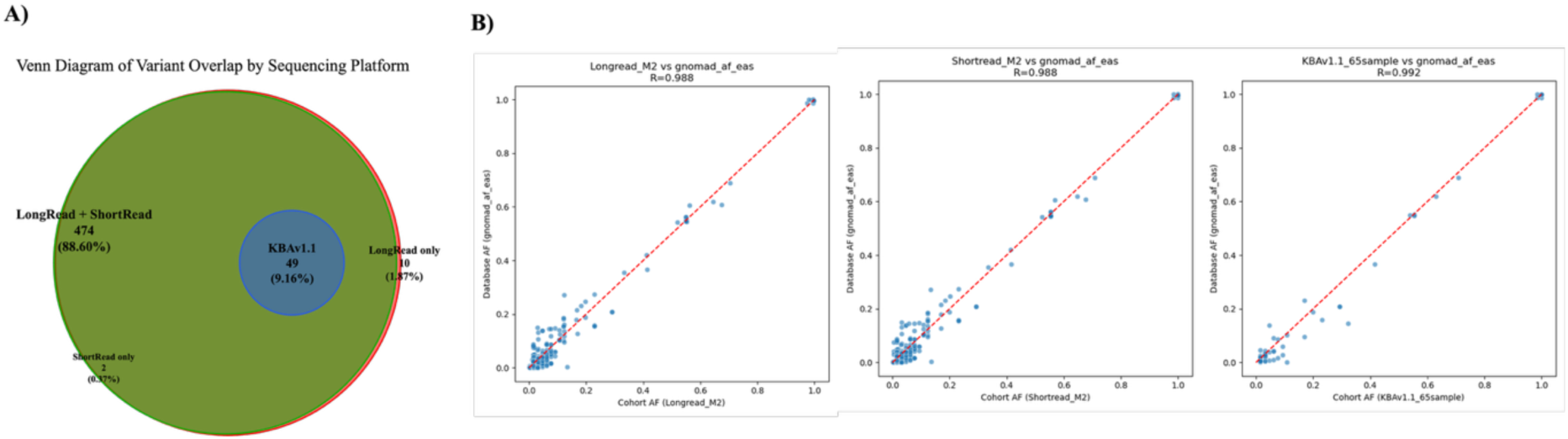
Cross-platform validation of variant calling and population genetic consistency. (a) Venn diagram of mitochondrial variants by sequencing platform. Analysis was restricted to polymorphic variants for comparison with SNP microarray data. Five variants with discordant genotypes between platforms were excluded. (b) Allele frequency correlation of common variants between gnomAD v3.0 EAS and the Korean dataset across sequencing platforms. Abbreviations: AF, allele frequency; GT, genotype; KBAv1.1, Korea biobank array version 1.1; R, Pearson correlation coefficient.

Variant-level confidence, as measured by QUAL scores (tumor log-odds; TLOD) from Mutect2 in mitochondrial mode, was robust across variant types. Known SNVs had a mean QUAL of 1,032.97 (n = 503), while known InDels had a mean QUAL of 394.41 (n = 28), consistent with the lower alignment certainty inherent to length-polymorphic loci but well above standard variant calling thresholds. The two novel variants showed lower but clearly detectable QUAL scores (76.24 for m.15635 T>A; 32.41 for m.4247 TT>T), reflecting their heteroplasmic nature rather than analytical uncertainty. Together, the high cohort allele frequency concordance with external databases and the robust variant-level QUAL distribution establish the analytical reliability of the LRS variant set.

Variant distribution across the mitochondrial genome reflected the expected mutational landscape: 403 variants (75.7%) mapped to the coding region (positions 577–16,023), comprising 389 SNVs and 14 InDels, while 130 variants (24.3%) mapped to the non-coding control region (D-loop and flanking sequences), comprising 115 SNVs and 15 InDels. Although the control region accounts for only ∼7% of the mitochondrial genome by length, it harbored 24.4% of all detected variants and 51.7% of all InDels, reflecting the well-established enrichment of length-polymorphic homopolymeric tracts and repetitive elements in the D-loop that are particularly susceptible to replication slippage.

To validate the discovered variants, we applied the identical Mutect2-based pipeline (Longread_M2) to the independent 1KGP Oxford Nanopore Technologies (ONT) long-read dataset (n = 1,019 individuals across five continental super-populations) (Schloissnig et al. 2025). The m.4247 TT>T deletion was independently detected in 42 of the 1,019 individuals at comparable heteroplasmy levels (mean VAF = 0.239; range: 0.118–0.330; mean QUAL/TLOD = 21.14), confirming it as a recurrent variant detectable by long-read platforms. In contrast, neither m.4247 TT>T nor m.15635 T>A was detected in the mitochondrial genome data from 1KGP Phase 3 high-coverage short-read dataset (n = 2,504; Illumina ≥30×) to which the Mutect2-based pipeline was applied (Byrska-Bishop et al. 2022). The absence of both variants from this large-scale short-read reference panel, together with their detection across two independent long-read cohorts, demonstrates that long-read sequencing reveals a class of mitochondrial variants systematically missed by conventional short-read approaches.

### Korean Mitochondrial Genomic Landscape through Haplogroup Analysis

Principal Component Analysis (PCA) using 1KGP Phase 3 reference populations confirmed that Korean individuals cluster within the East Asian (EAS) super-population, clearly separated from African (AFR), European (EUR), South Asian (SAS), and admixed American (AMR) groups along PC1 and PC2 (**Figure 3b, 3c**). Within the EAS cluster, Korean samples showed partial overlap with Han Chinese (CHB, CHS) and Japanese (JPT) populations but occupied a slightly distinct position, reflecting population-specific haplogroup frequency differences consistent with Northeast Asian population history.

**Figure 3.**
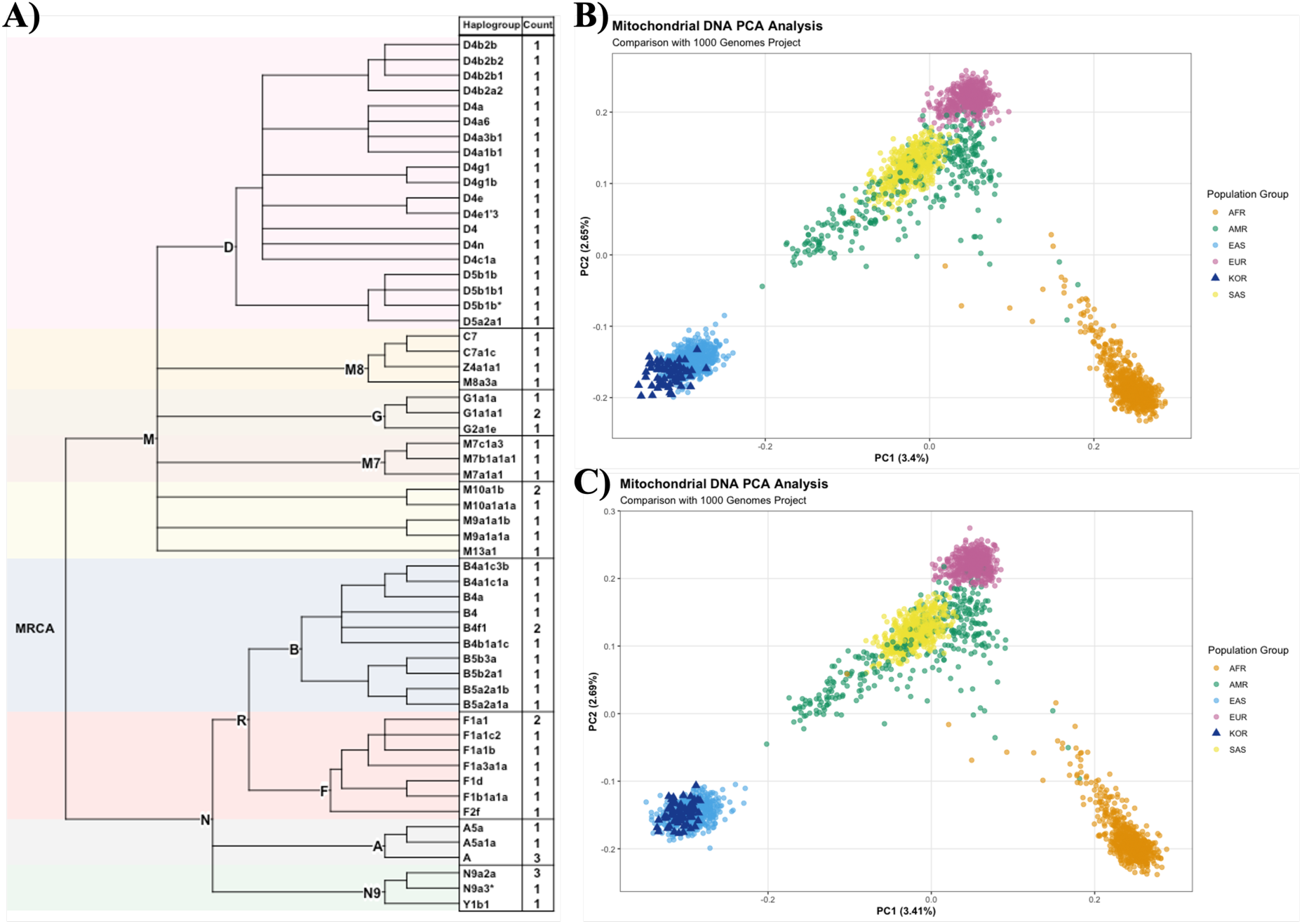
Mitochondrial haplogroup analysis and population structure of 65 Korean individuals. (a) Frequency distribution of mitochondrial haplogroups in the Korean population. (b) Principal Component Analysis (PCA) plot comparing 65 Korean individuals long-read sequencing with samples from the 1000 Genomes Project Phase 3. (c) PCA plot comparing 65 Korean individuals short-read sequencing with samples from the 1000 Genomes Project Phase 3.. Abbreviations: MRCA, most recant common ancestor; AFR, African; EUR, European; SAS, South Asian; EAS, East Asian; AMR, Amerian; CHS, Han Chinese South; CHB Han Chinese Beiging; JPT, Japnese in Tokyo; POS, position; REF, reference allele; ALT, alternative allele; GT, genotype; VAF, variant allele frequency; DP, depth; TLD, tumor log odds.

Using the LRS-derived variant set, we characterized the mitochondrial genomic profile of the 65 Korean individuals. Haplogroups were assigned using HaploGrep3 based on PhyloTree Build 17 (Forensic Update 1.2). The cohort displayed a haplogroup distribution characteristic of East Asian populations: macro-haplogroup D was the most prevalent (29.2%; n = 19), comprising D4 (23.1%; n = 15) and D5 (6.2%; n = 4), followed by macro-haplogroup B (16.9%; n = 11; B4 = 10.8%, B5 = 6.2%), macro-haplogroup M excluding C and Z (15.4%; n = 10; M7 = 4.6%, M8 = 1.5%, M9 = 3.1%, M10 = 4.6%, other M = 1.5%), F (12.3%; n = 8), A (7.7%; n = 5), G (6.2%; n = 4), N9 (6.2%; n = 4), C (3.1%; n = 2), Y (1.5%; n = 1), and Z (1.5%; n = 1) (**Figure 3a**).

The mtDNA haplogroup distribution in our cohort (n = 65) showed strong clade-level concordance with previously published Korean studies, with D4 predominating at 25–30%, macro-haplogroup B accounting for 10–15%, M sub-lineages collectively representing 15–20%, and haplogroups A, F, and G each contributing approximately 5–10% (Lee et al. 2006; Jin et al. 2009; Park et al. 2017). Both northern Siberian-affiliated lineages (D4, G, Y, Z) and southern East Asian-affiliated lineages (B, F, M7) were present at substantial frequencies, recapitulating the characteristic North–South admixture signature of Korean mtDNA. These results confirm that, despite its modest size, the cohort captures the major features of Korean maternal lineage diversity; moreover, the enhanced sensitivity of LRS—particularly for InDels in homopolymeric regions—is expected to reveal lineage-specific signals that have been inaccessible through prior short-read or microarray approaches.

### Cross-Platform Evaluation of Variant Detection

To better understand enhanced utility of LRS over existing platforms, we performed systematic cross-platform comparison using the same 65 individuals sequenced with short-read sequencing (SRS; Illumina NovaSeq, 150 bp paired-end; mean mitochondrial depth: 3,530.65×) and genotyped on the Korea Biobank Array v1.1 (KBAv1.1; 180 mitochondrial SNP probes). SRS detected 525 variants (504 SNVs, 21 InDels), while KBAv1.1 identified 54 non-monomorphic variants (**Table 1**). Concordance analysis revealed that 523 variants were called by both LRS and SRS, representing 98.1% of the LRS variant set (**Figure 2a**). Among these shared variants, 49 overlapped with KBAv1.1 calls (49 of 54 non-monomorphic KBA probes). Although platform specific variants were identified, most exhibited VAFs near the baseline filtering threshold (10%) indicating that they represent low-level heteroplasmic variants near the limit of platform sensitivity rather than technical errors in variant calling process **(Supplementary Table 2)**.

Variant-level overlap was highly concordant between platforms [SRS vs. LRS, 99.62%, n = 523 overlapping variants] and showed strong agreement with the Korean Biobank Array [KBA vs. LRS, 100.00%, n = 49 overlapping variants]. The inter-platform agreement indicates that SRS and LRS yield equivalent population-level allele frequency estimates, while the concordance with KBA further supports that neither platform deviates systematically from an orthogonally genotyped Korean cohort.

Although SRS achieved approximately 14-fold higher mean sequencing depth than LRS (3,530.66× vs. 254.81×) and correspondingly higher per-variant statistical confidence (mean QUAL / TLOD of 12,144.54 for SRS known variants vs. 1,006.21 for LRS known variants), this depth advantage did not translate into greater variant discovery sensitivity. On the contrary, LRS identified 38.1% more InDels than SRS (29 vs. 21) despite substantially lower depth (**Table 1**), indicating that sequencing depth and variant detection breadth are partially decoupled for the mitochondrial genome: the continuous read architecture of LRS provides a distinct analytical advantage that per-variant statistical confidence alone does not capture.

To evaluate whether the variant-level differences observed between platforms translate into measurable consequences for downstream analysis, we compared haplogroup assignment accuracy across LRS, SRS, and KBAv1.1 by paired testing of phylogenetic depth, quality scores, and missing haplogroup-defining variants (**Table 2**). No significant differences were observed between LRS and SRS (mean haplogroup depth: 13.52 vs. 13.54 levels, p = 0.321; quality score: 0.970 vs. 0.971, p = 0.153; missing variants: 0.169 vs. 0.200, p = 0.321), consistent with their equivalent variant-level concordance. However, the haplogroup A-specific novel InDel **<u>(chrM:4247:TT,T)</u>** could only be recovered by LRS, further illustrating how its analytical advantages propagate into biologically meaningful downstream findings.

**Table 2.** Paired t test for statistic of mitochondrial genome haplogroup.

| Sequencing Platform | Haplogroup Depth<br>(PhyloTree Build 17-FU 1.2) |  |  | Quality Score<br>(Haplogrep3) |  |  | Number of missing<br>variants for Haplogroup |  |  |  |
| --- | --- | --- | --- | --- | --- | --- | --- | --- | --- | --- |
|  | Mean | SD | *p* value | Mean | SD | *p* value | Number | Mean | SD | *p* value |
| Longread | 13.523 | 2.144 | Ref | 0.970 | 0.025 | Ref | 11 | 0.169 | 0.417 | Ref |
| Shortread | 13.538 | 2.166 | 0.3211 | 0.971 | 0.025 | 0.1529 | 13 | 0.200 | 0.474 | 0.3211 |
| KBAv1.1 | 11.400 | 2.082 | 4.4803E-11* | 0.979 | 0.029 | 0.0314* | 5 | 0.077 | 0.269 | 0.1347 |

\**p* < 0.05
Abbreviations: PhyloTree Build 17-FU 1.2, PhyloTree Build 17-Forensic update 1.2; SD, standard deviation.

KBAv1.1, however, achieved significantly lower phylogenetic depth (mean 11.40 levels; p = 4.48 × 10⁻¹¹ vs. LRS) despite a marginally higher nominal quality score (0.979; p = 0.031). This pattern reflects the design of KBAv1.1: its probe set targets canonical haplogroup-defining SNPs, yielding high accuracy at the macro-haplogroup level but lacking resolution for fine-scale sub-haplogroup classification that requires comprehensive sequence data. The approximately two-level reduction in phylogenetic depth with KBAv1.1 represents a meaningful loss of resolution for studies requiring sub-haplogroup-level maternal lineage characterization—particularly in population genetic contexts where fine-scale haplogroup sub-structure carries phylogeographic significance.

### Functional Characterization of Novel Variants

#### *MT-ND1* Frameshift Variant (m.4247 TT>T; p.Ile314X)

This single-base deletion in a TT dinucleotide creates a frameshift at codon 314 of ND1 (p.Ile314X), altering the amino acid sequences at the C-terminus. The variant was called as a heteroplasmic allele (VAF = 0.103) in one individual assigned to Haplogroup A and was detected exclusively by LRS; no SRS call was made at this position despite a mean mitochondrial depth of 3,530.63× (**Table 3)**.

**Table 3.**
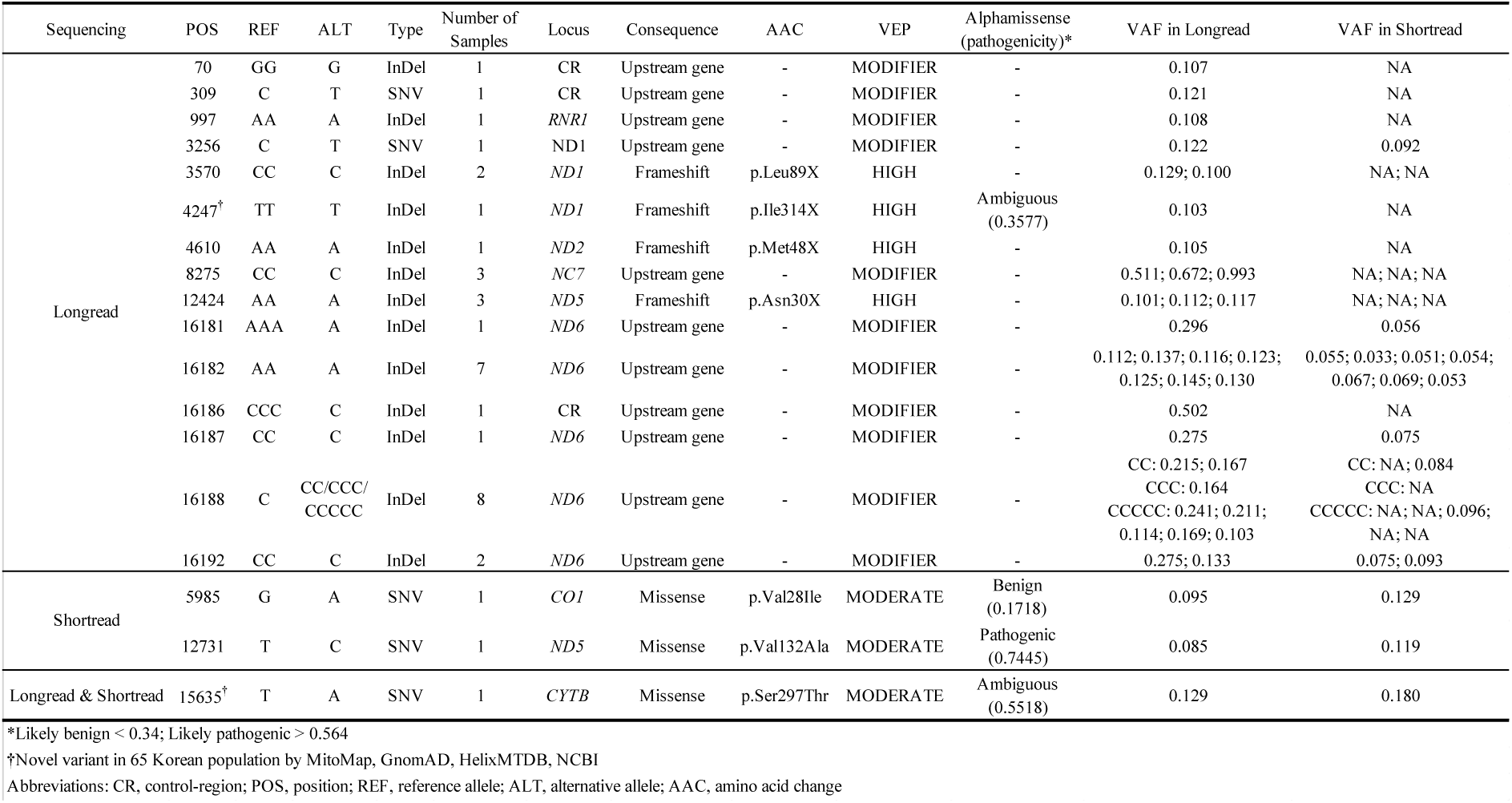
List of unique variants by sequencing platform.

Haplogroup A comprised five individuals in our cohort, all of whom carried the canonical motif variant m.4248 T>C as a homoplasmic substitution (VAF 0.895–0.948). LRS additionally detected the adjacent frameshift deletion m.4247 TT>T in all five individuals: one above the heteroplasmy calling threshold (VAF = 0.103) and four at sub-threshold levels (VAF 0.052–0.061; mean depth 154×) visible by direct inspection of aligned long reads. In the 1KGP ONT dataset (Schloissnig et al., 2025), the same variant was recovered in East Asian and admixed American samples carrying Haplogroup A lineages and was not observed in other haplogroups (**Supplementary Table S1**).

AlphaFold3 modeling of the protein structure of frameshift mutation (m.4247 TT>T) showed loss of the C-terminal region containing Thr164 (mature numbering) and of the associated hydrogen-bond contacts with adjacent Complex I subunits (**Figure 4A**). FoldX returned ΔΔG = −45.21 kcal/mol (**Supplementary Figure 1**). A 100-ns GROMACS MD simulation in a model mitochondrial inner membrane showed divergent trajectories: the wild-type remained stable (RMSD < 1.5 Å), whereas the mutant exhibited decreased Radius of Gyration and SASA, altered C-terminal RMSF, altered hydrogen bonding, and RMSD > 2.0 Å relative to the equilibrated wild type (**Supplementary Figure 2**). These features are consistent with compaction of the substituted C-terminus accompanied by its detachment from neighboring subunits, and over the simulated timescale the variant failed to sample the wild-type-like hydration-coupled open conformation, suggesting a reduced propensity for the proton-gating transition of Complex I. These effects could reduce the proton motive force and ATP-generating capacity, potentially increasing weakness in tissues with high energy demands, such as the optic nerve (Carelli et al. 2004; Hirst 2013; Maresca et al. 2013). Consistent with this possibility, pathogenic variants in *MT-ND1* have been previously associated with Leber hereditary optic neuropathy (LHON) (Ji et al. 2019).

**Figure 4.**
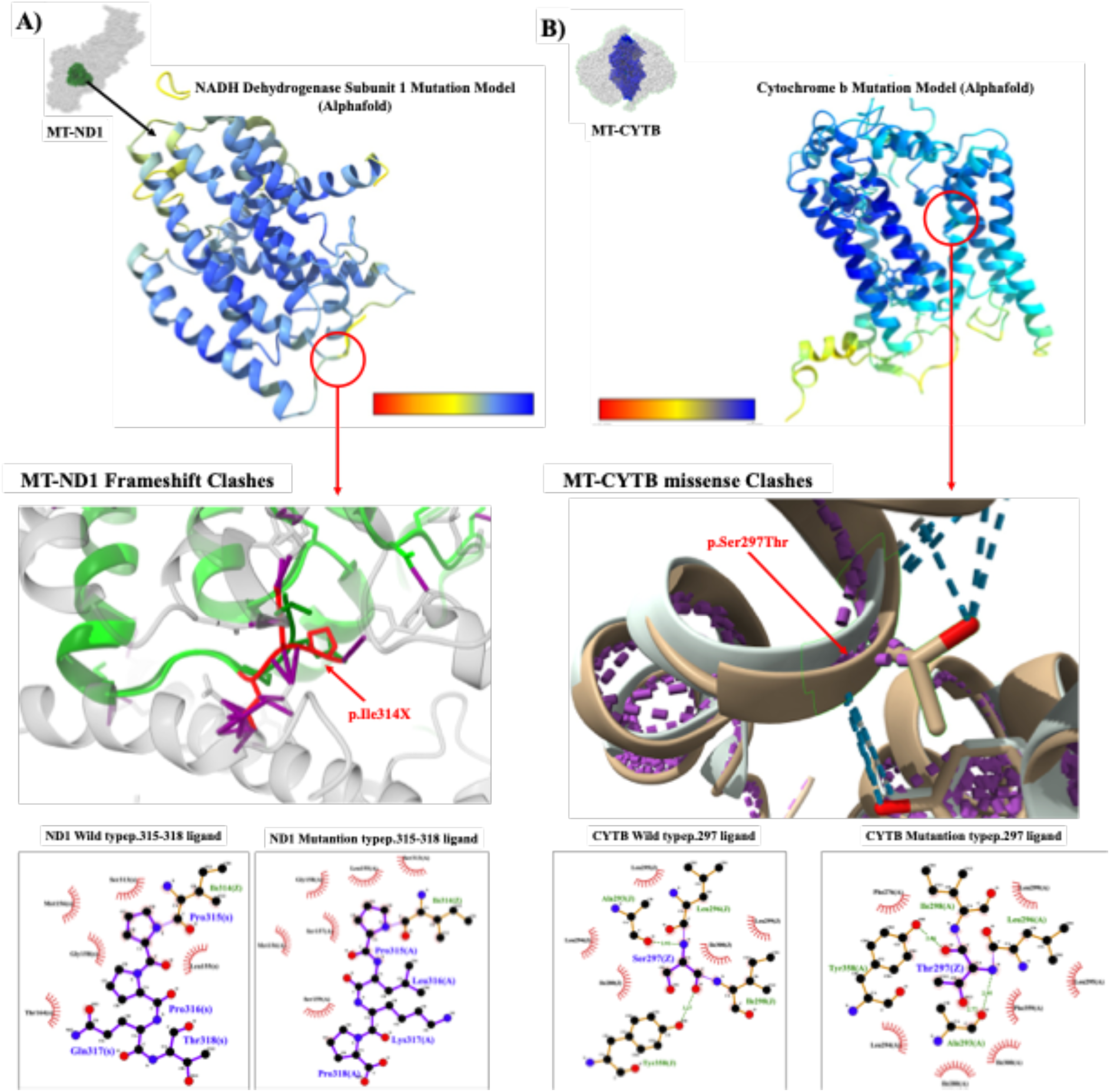
Structural modeling and effect prediction of the ND1 protein with the m.4247 TT>T variant and CYTB protein with the m.15635 T>A variant. (A) ND1 protein structure modeled by AlphaFold 3 based on the m.4247 TT>T sequence. (B) CYTB protein structure modeled by AlphaFold 3 based on the m.15635 T>A sequence. Abbreviations: WT, wild type.

*MT-CYB* Missense Variant (m.15635 T>A; p.Ser297Thr)

This single-nucleotide substitution introduces a Ser→Thr replacement at codon 297 of cytochrome b. The variant was called as a heteroplasmic allele in one individual assigned to Haplogroup M10a1b and was detected by both platforms LRS (VAF = 0.129; DP = 199) and SRS (VAF = 0.180; DP = 3,930). AlphaMissense assigned an “Ambiguous” score (0.5518) and VEP a MODERATE impact (**Table 3**).

AlphaFold3 modeling of the substituted protein showed a preserved overall fold (RMSD < 1.0 Å) with localized rearrangement at position 297 (**Figure 4B**). FoldX position scanning returned ΔΔG ≈ +0.94 kcal/mol for the Ser→Thr substitution. LigPlot+ analysis showed altered intermolecular contacts between the Thr297 side chain and the bound ubiquinone at the Qo oxidation site (**Figure 4B**). This alteration may affect Quinone cycle dependent electron transport chain in Complex III, therefore reducing respiratory-chain efficiency (Andreu et al. 1999; Crofts 2004). Although another substitution at the same residue, p.Ser297Pro, has been associated with complex III deficiency, m.15635 T>A showed discordant *in silico* predictions and modest effects (VEP: MODERATE; AlphaMissense: Ambiguous) (**Table 3)** (Fragaki et al. 2009; Tropeano et al. 2025). Therefore, this variant is interpreted as a uncertain functional significance with plausible local effects on Complex III.

## Discussion

This study demonstrates that long-read sequencing provides a more complete view of human mitochondrial genome variation than conventional short-read sequencing or genotyping arrays. In 65 Korean individuals, LRS generated the most abundant mtDNA variant catalog despite lower sequencing depth, while maintaining high concordance with SRS for single-nucleotide variants and haplogroup assignment. This concordance establishes LRS as a reliable baseline technology, but the central finding is that LRS recovered variants that were missed by SRS and arrays, particularly insertion-deletion events in regions that are difficult to resolve with short reads. Notably, the haplogroup A-associated MT-ND1 InDel m.4247 TT>T was detected by LRS and independently replicated in 1KGP ONT data, supporting its authenticity rather than a platform-specific artifact. These findings indicate that the apparent completeness of current mtDNA variant catalogs is partly constrained by the technologies used to build them, and they argue for systematic incorporation of LRS-derived mitochondrial variants into population and clinical resources such as gnomAD and ClinVar.

The key strength of this work lies in what mitochondrial genome analysis based on LRS reveals about an underrepresented population. By resolving full-length mitochondrial molecules, LRS discovered homopolymer-associated InDels and lineage-specific variants in the Korean. This population are one of the insufficiently represented groups in mitochondrial reference resources, comprising only a small fraction of East Asian samples in gnomAD which conventional short-read and array databases leave unresolved. This matters because such variants are precisely those excluded from rCRS and SRS based databases, so their recovery directly features a reference-bias gap rather than re-detecting known polymorphisms. The credibility of this claim rests on three design features. First, the same 65 individuals were assayed on all three platforms, removing the cohort and sampling confounders that limit most benchmarking studies (Foox et al. 2021). Second, the headline LRS-specific finding was reproduced in an independent cohort based on LRS technology (1KGP ONT). Third, connected to population-genetic structure and protein structural modeling, moving the result from a variant catalog toward a biologically interpretable observation.

Among the novel variants, the *MT-ND1* frameshift deletion (m.4247 TT>T; p.Ile314X) is the most functionally compelling. This variant implied a loss of inter-subunit hydrogen bonds, reduced solvent exposure. These features may cause potential disturbances of Complex I proton-gating dynamics. Because ND1 is a core Complex I subunit and *MT-ND1* variation has repeatedly been reported associations to OXPHOS phenotypes, including low-penetrance LHON modification and isolated Complex I deficiency, while the immediately adjacent haplogroup A motif m.4248 T>C lies in the same functional neighborhood (Ji et al. 2019; Ng et al. 2020). At the cellular level, impaired Complex I proton-gating may reduce respiratory reserve and ATP-generating capacity while increasing redox stress (Hirst 2013; Sazanov 2015; Hirst and Roessler 2016). At the tissue and organism levels, such effects would be expected to preferentially affect high-energy tissues, including neural, optic, muscular, and cardiac systems (Carelli et al. 2004; Maresca et al. 2013; Ji et al. 2019; Ng et al. 2020). Nevertheless, the variant was heteroplasmic and no *in vitro or vivo* validation was performed, so it should be regarded as a high-priority candidate functional variant rather than a confirmed pathogenic mutation, with its haplogroup A recurrence equally consistent with lineage-specific tolerance or drift.

The *MT-CYB* variant (m.15635 T>A; p.Ser297Thr) needs more conservative reading. It was detected concordantly by LRS and SRS in a single haplogroup M10a1b individual, but it is a clinically uncertain variant. Although a different substitution at the same residue, p.Ser297Pro, has been reported in severe Complex III disease, this same-site precedent indicates that Ser297 is functionally sensitive rather than that p.Ser297Thr is itself pathogenic (Fragaki et al. 2009; Tropeano et al. 2025). *MT-CYB* encodes cytochrome b, the mtDNA-encoded catalytic core subunit of respiratory Complex III (Tucker et al. 2013; Fernández-Vizarra and Zeviani 2015). Therefore, alteration of a conserved cytochrome b residue could affect ubiquinol-cytochrome c electron transfer and Q-cycle-dependent proton-gradient generation (Iwata et al. 1998; Crofts 2004). Such disruption of Complex III function may reduce respiratory-chain efficiency and ATP-generating capacity and may alter mitochondrial redox balance (Andreu et al. 1999; Fragaki et al. 2009; Ekiert et al. 2016; Tropeano et al. 2025). Therefore, we suggest that functional follow-up study is necessary for this.

Taken together, these findings clarify why read length, and not depth alone, affects the breadth of mitochondrial variation we found. The mitochondrial genome is a compact, homopolymer-rich circular molecule mirrored by hundreds of nuclear-mitochondrial segments (NUMTs), so short reads remain error-prone there even at very high depth, whereas full-length long reads span these regions in single molecules and discriminate true mtDNA from NUMT-derived signal (Wei et al. 2022; Macken et al. 2023). This is why greater SRS depth did not translate into broader discovery, and we have confirmed that LRS can be used as a high-resolution method for mitochondrial analysis. But SRS remains highly accurate and scalable for shared SNVs and population-scale studies, and arrays, which remain cost-effective for macro-haplogroup inference. Practically, LRS can serve as a reference-grade approach for discovering and curating complex mitochondrial variation and incorporating long-read-resolved variants into public databases, phylogenetic trees, and future population-specific mitochondrial pangenome resources will be central to reducing reference bias for underrepresented Korean and broader East Asian populations (Laricchia et al. 2022; Liao et al. 2023; Schloissnig et al. 2025).

This study has several limitations. First, the Korean cohort was modest in size, so rare-variant frequencies and haplogroup-specific enrichment should be treated as carefully; the matched three-platform design nonetheless provides high internal validity for the platform comparison. Second, functional interpretation of the novel variants rests on *in silico* modeling and molecular dynamics simulation, which generate mechanistic hypotheses but cannot replace experimental validation. Third, heteroplasmic variants near the 10% VAF threshold remain challenging for all platforms, and sub-threshold observations require cautious interpretation. Finally, current mitochondrial phylogenetic frameworks and databases predate long-read resequencing and remain incomplete for population-specific InDels, so some LRS-detected variants may appear “novel” less because they are exceptionally rare than because prior technologies were not optimized to capture them.

In summary, this study demonstrates that LRS provides added resolution for mitochondrial genome analysis, particularly for InDel discovery and lineage-specific variants that are difficult to recover with short reads or arrays. The findings support the use of LRS as a complementary reference-level approach for mitochondrial variant discovery, database refinement, and the future construction of population-aware mitochondrial pangenome resources.

## Acknowledgments

This work was supported by an intramural grant from the Korea National Institute of Health (2024-NI-007-02). This study was performed with bioresources from National Biobank of Korea, the Centers for Disease Control and Prevention, Republic of Korea. Informed consent was obtained from the participants involved (Approved IRB number of Korea Centers for Disease Control and Prevention in Korea: KDCA-2024-02-09).

## Data Access

Long-read sequencing data, including allele-frequency information in VCF format, are publicly available on the KPP GitHub repository (https://github.com/KoreanPangenome/KPPD).

## Ethics statement

All participants provided written informed consent. This study was approved by the Institutional Review Board of the Korea Disease Control and Prevention Agency, Republic of Korea (IRB number: KDCA-2024-02-09).

## Author contributions

B.-J.K., H.J.J., and Y.J.K. contributed to the design of the study. H.J.K., B.-J.K., H.J.J., and Y.J.K. wrote the manuscript. H.J.K., S.M.K., and K.Y. performed data curation, analysis, and validation. All authors interpreted the results. All authors approved the submission of the final version of the article for publication.

